# A Method for Image-Based Modeling of Uterine Passive Mechanics During Late Pregnancy

**DOI:** 10.64898/2026.07.10.737823

**Authors:** Olivia Mergler, Abigail Laughlin, Erin Louwagie, Lei Shi, Kristin Myers, Vijay Vedula

## Abstract

**Purpose:** Computational models of the uterus during pregnancy enable analysis of electro-chemo-mechanical pathways to predict labor timing and guide treatment planning. We aim to develop a robust image-based modeling pipeline to investigate uterine passive mechanics during late pregnancy.

**Methods:** A parametric model of the uterus and cervix was created using a patient’s MRI measurements at 38 weeks of gestation. Inspired by advances in cardiac mechanics models, we created Laplace-Dirichlet solutions to inform tissue domains, fiber structure within the uterus and cervix, and spatially varying Robin boundary conditions. Prior imaging and mechanical testing data were used to fit material parameters. Boundary condition parameters were tuned to match the displacements of a previously established approach that employed contact with surrounding tissue. The tissue mechanical response to a physiologic load was assessed across varying material properties and fiber architectures.

**Results:** Discrepancies in nodal displacements between the current approach and the contact-based model were limited to 3.4 ***±*** 1.8 mm, yielding nearly 90 % computational savings. Uterine tensile strains were more sensitive to ground substance elastic modulus (***E***) compared to fiber properties. Reduced ***E*** and fiber stiffness increased cervical strains and compression. Fiber dispersion and architecture modulated the opening of the cervical internal ostium but had a reduced impact on compression.

**Conclusion:** We developed a novel workflow for modeling passive uterine mechanics, informed by patient-specific measurements and *in vitro* mechanical tests. The robust workflow may prove useful for studying labor progression and conducting longitudinal studies to enhance our understanding of normal and pathological pregnancies.

## 1 Introduction

Approximately 3.6 million pregnancies occur in the United States annually [1]. Yet we do not have an accurate predictive mechanism for determining when a pregnant mother will enter labor. The estimated delivery date (EDD) is approximately 280 days from the last menstrual period and is further refined using ultrasound imaging during early gestation [2]. However, only 5 % of pregnant women actually enter labor on the EDD, with about 34 % of births occurring at least a week outside of the EDD (± 7 days) [2]. Poor prediction in normal pregnancies makes it even more challenging to predict the course of abnormal pregnancies, such as preterm birth or post-term birth [3]. Preterm birth, defined as birth before 37 weeks of gestation, accounts for 10 % of births worldwide, a rate that has continued to rise over the past decade [4]. Due to insufficient fetal development at this stage, preterm birth is the leading cause of childhood mortality, as well as lasting childhood disability, including neuro-developmental, respiratory, and cardiovascular complications [4, 5]. Alternatively, post-term birth is defined as a pregnancy that extends beyond 42 weeks of gestation, accounting for 7 % of all pregnancies [6]. Post-term births increase the risk of fetal distress during labor, neonatal seizures, perinatal mortality, requirement of a C-section, and postpartum maternal hemorrhage [7].

Mechanical processes play a crucial role during pregnancy and the onset of labor. Throughout pregnancy, the uterus grows and distends to accommodate fetal growth [8]. The cervix functions as a mechanical barrier, preventing the opening of the uterus before labor, and prepares for labor by softening (decreased collagen crosslinking and mechanical stiffness) and dilating at the end of the pregnancy [8–10]. In addition to hormonal regulation, the growth and remodeling of both tissues are largely driven by stretch-mediated inflammatory pathways [9]. For example, fetal membrane distension has been shown to increase the presence of the proinflammatory cytokine IL-8 and collagenase activity, two mediators of cervical ripening [11]. Such a phenotype is necessary for labor onset but can contribute to preterm labor if it occurs too early [12]. Alternatively, reduced IL-8 levels were found in post-term labor patients who did not respond to labor induction and ultimately required a C-section delivery, compared to those who experienced cervical ripening upon labor induction [7]. Further, throughout pregnancy, the uterus remains quiescent and becomes highly excitable just before labor, a transition mediated by inflammatory processes [13, 14]. The presence of stretch-activated ion channels, particularly PIEZO channels, has been found to be a critical regulator of myometrial^1^ excitability by facilitating cell depolarization under mechanical stretch [16, 17]. Therefore, uterine stretch is a key mediator of labor onset, and uterine overdistension has been found to contribute to preterm labor [9, 18].

Understanding the mechanical environment that modulates these mechanosensitive regulatory pathways may help predict the onset of labor in both physiological and pathological pregnancies. Image-based computational models offer a high-fidelity, noninvasive means to investigating the passive mechanics of the uterus and cervix, providing a baseline understanding of the mechanical signals that arise under physiological loading. When placed in the context of the biomechanical pathways that drive pregnancy and labor progression, such studies can provide insight into how different mechanical environments may affect pregnancy outcomes [19]. These models can be further advanced to synthesize known biomechanical pathways and analyze the sensitivities that differentiate physiological and pathological labor conditions, including pre- and post-term births [20–22]. The use of such models enable the development of patient-specific approaches to guide clinical treatment. By advancing predictive capabilities for pregnancy, we can assess and optimize the timing and choice of early intervention strategies for pathological pregnancies, such as mechanical interventions (e.g., cerclage) or drug delivery to modulate hormone regulation [23–25]. Such models can also be used to understand risks during childbirth to both the mother and fetus [26, 27]. In multiparous cases, computational modeling could be used to predict the biomechanical response of the uterine scar tissue following a prior Cesarean delivery [28].

Several computational models have been developed to characterize the biomechanical response of the uterus under passive physiological loads, as well as during active contractions just before or during labor [21, 27, 29–32]. Such models are employed to understand the sensitivity of the mechanical response to different tissue geometries, material properties, and excitation-contraction dynamics. Previous models of uterine mechanics have relied on simplified boundary conditions, such as a fixed base of the uterus [29, 33, 34], or physiologically-based approaches, where the uterus is bounded by external tissue components, including the abdomen, pelvis, or pelvic floor muscles [19, 21, 22, 35].

This work aims to build upon these previous approaches, improving the predictive value of uterine mechanical modeling through the development of a computationally efficient, yet physiologically-informed model configuration. Such an approach will ultimately reduce computational barriers when scaled across gestation for longitudinal studies that account for uterine growth and remodeling and active contraction during labor.

Within the fields of cardiac and vascular biomechanics, these aims have been routinely approached, and such developments can be translated to uterine biomechanics applications to improve model robustness and efficiency. For the heart, motion constraints often come from the pericardium and other organs of the thoracic cavity, such as the diaphragm, lungs, and rib cage [36–38]. For the aorta, there are interactions with external tissues, including the heart, the spine, and the attached branch vessels and intercostal arteries [39–41]. For the uterus, motion is constrained by the fetal membrane and organs throughout the upper body, such as the bowel, spine, pelvis, and pelvic floor muscles, as well as attachment to ligaments [35, 42]. Robin-type spring-dashpot systems have been adopted as boundary conditions to effectively mimic these motion constraints in cardiovascular mechanics applications [36, 38, 39, 41, 43, 44]; thus, they could potentially be applied to constrain uterine motion. Furthermore, both cardiovascular tissues and the uterus exhibit nonlinear stress-strain relations, anisotropy, and tension-compression asymmetry, motivating the development of anisotropic constitutive models for both tissues to characterize a fiber-embedded material that can be used in passive mechanics studies [19, 45, 46].

In this work, we aim to develop a novel and robust image-based uterine mechanics modeling workflow, specifically designed to facilitate seamless future integration into contraction and longitudinal studies. The current study will focus on the initial model configuration, verification, and investigation of the passive uterine mechanics under physiological loading during late pregnancy. Established approaches to modeling cardiovascular biomechanics will serve as the basis for each of the steps implemented here, including fiber generation techniques, material model selection, and boundary condition application, thereby enabling a balance between biofidelity, model robustness, and efficiency.

## 2 Materials and Methods

### 2.1 Modeling Workflow

We developed a novel workflow for image-based modeling of uterine passive mechanics, leveraging advances in cardiac biomechanics. Major components of this workflow include: (i) generating an image-based geometry from a patient’s data (Section 2.1.1), (ii) creating Laplace solutions with Dirichlet boundary conditions (Section 2.1.2), which then informs (iii) uterus subdomains (Section 2.1.3), (iv) fiber generation (Section 2.1.4), and (v) boundary condition development using spatially varying Robin boundary conditions (Section 2.1.7). Additional key components include material characterization using an anisotropic, hyperelastic constitutive model (Section 2.1.5) and parameter estimation (Section 2.1.6). Each of these steps will be discussed in detail in the following subsections.

#### 2.1.1 Image-Based Geometry

We have adopted a previously developed parametric modeling pipeline to create a three-dimensional geometric model of the uterus during late pregnancy from patient imaging (Fig. 1) [22]. The subject (Subject 05) used in this model was imaged at 38 weeks’ gestation using magnetic resonance imaging (MRI) [22]. We note that the parametric modeling approach employed here has also been implemented using measurements derived from two-dimensional ultrasound imaging [19, 47]. While the present study relied on MRI data, the ability of this method to use routinely acquired ultrasound data enables scalability across a broader population and minimizes barriers to clinical translation [22, 48]. Briefly, this parametric modeling approach employed imaging data to derive patient-specific dimensions of uterine and cervical geometries, including diameters, wall thicknesses, and the angle of the cervix relative to the uterus [49] (Fig. 1, left). In SolidWorks (Dassault Syst*è*mes, Inc., Version 2025 Education Edition, Vélizy-Villacoublay, France), these measurements were used to parameterize a generic template for uterine and cervical geometry. For the uterine geometry, a series of ellipses and splines that made up the inner and outer uterine wall outlines were dimensioned based on sagittal plane geometries. This geometry was then lofted based on elliptical profiles defined in the coronal plane. The cervix consists of an extruded cylinder, smoothed with fillets [19].

**Fig. 1.**
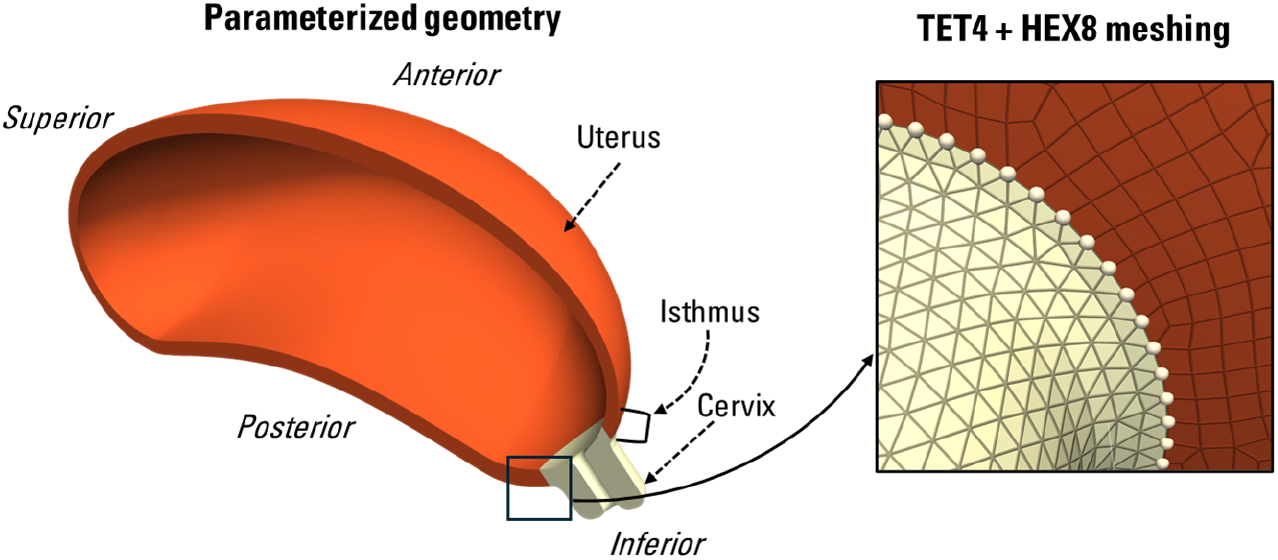
Image-based finite element model construction of a pregnant uterus. (left) Parameterized uterine geometry for a subject at 38 weeks of gestation with anatomical directions and pertinent tissue domains labeled. (right) A hybrid mesh is used for the uterus and cervix, with the cervix meshed using four-noded linear tetrahedral elements (TET4) and the uterus using eight-noded trilinear hexahedral brick elements (HEX8) with node-matched continuity between the two domains at the interface.

Once the parameterized geometry was built, we then meshed the tissue volume for subsequent finite element analysis (FEA) (Fig. 1). To optimize mesh density across the long, slender uterus and the relatively short, thicker cervix, we adopted a hybrid meshing strategy. The cervix was meshed with linear tetrahedral elements (TET4) ranging from 0.4 mm on the inner canal to 2 mm on the outer surface. The uterus was meshed with trilinear hexahedral elements (HEX8) using solid map meshing in Hypermesh (Altair, Inc., Version 2022, Troy, MI). The hexahedral elements have an edge length of 2 mm in the in-plane direction and a depth ranging from 1 to 2.5 mm, enabling five elements to span the thickness of the entire uterus. All mesh sizes were chosen based on mesh size convergence studies performed in Louwagie (2024) [19]. The meshes of each subdomain (i.e., the uterus and cervix) share attributes, including nodes, edges, and faces at the interface, creating a continuous mesh throughout the tissue and eliminating the need for contact application (Fig. 1, right). The resulting image-based parameterized FEA model consists of 89,825 HEX8 elements in the uterus and 69,903 TET4 elements in the cervix (159,728 elements in total).

#### 2.1.2 Laplace-Dirichlet Solutions

In our workflow, several model configuration steps, including the definition of tissue subdomains, fiber generation, and spatially varying boundary conditions, relied on Laplace-Dirichlet solutions [50, 51]. Each of these solutions was created by solving the Laplace equation for a scalar field (∇^2^*ϕ* = 0) over a domain of interest with appropriately chosen Dirichlet boundary conditions (Fig. 2a). Four different Laplace solutions were pertinent to this work: (i) a transmural solution from the outer surface to the inner surface of the uterus and cervix, *ϕ*_*TM*_, (ii) a full-geometry apico-basal solution, from the apex of the uterus (fundus) to the base of the cervix facing the vaginal canal, *ϕ*_*AB*_, (iii) an anterior-posterior solution from the anterior to the posterior surface of the uterus (cervix included in solution), *ϕ*_*AP*_, and (iv) a uterine apico-basal solution from the apex of the uterus to the uterus-cervix interface, *ϕ*_*AI*_ (Fig. 2a).

**Fig. 2.**
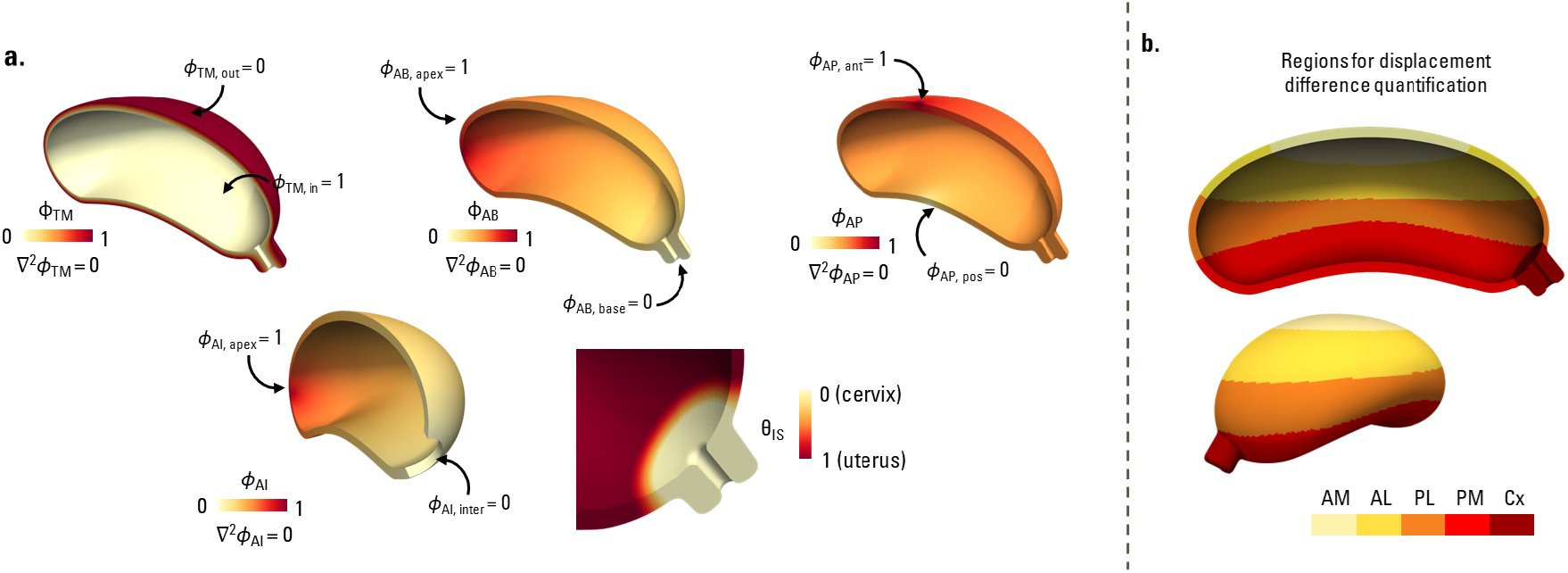
Laplace-Dirichlet parameterization to inform material subdomains and fiber directions. (a) Laplace solutions with Dirichlet boundary conditions are used for domain assignment, fiber generation, and boundary condition development. (top-left) *ϕ*_*TM*_ defined in the transmural direction, (top-center) *ϕ*_*AB*_ in the apico-basal direction, (top-right) *ϕ*_*AP*_ in the anterior-posterior direction, and (bottom-left) *ϕ*_*AI*_ defined in the apico-basal direction for the uterus alone. (bottom-right) *ϕ*_*AI*_ was mapped using *θ*_*IS*_ to define the isthmus layer, enabling a smooth transition of material properties and fiber directions between the cervix and the uterus. (b) *ϕ*_*AP*_ was used to define regions across the uterus, facilitating a quantitative comparison of the model-predicted deformation (Table 5), viewed along (top) the sagittal plane and (bottom) an oblique plane. AM: antero-medial; AL: antero-lateral; PL: postero-lateral; PM: postero-medial; Cx: cervix.

#### 2.1.3 Material Subdomains

The parametric modeling workflow in Section 2.1.1 resulted in creating two subdomains for the uterus and cervix. However, morphometry and immunohistological studies identified the presence of an isthmus region between the uterus and cervix, serving as a mechanical transition region between the different tissues [52, 53]. This region has been approximated to range from 6 to 10 mm in the nonpregnant uterus, extending radially from the cervix [54]. We incorporated this isthmus layer into our model to facilitate a smooth transition of the material parameters between the uterus and cervix. A clamped linear function, *θ*_*IS*_, was defined mapping the scalar field *ϕ*_*AI*_ to define the isthmus layer as an additional subdomain of the uterus, as,

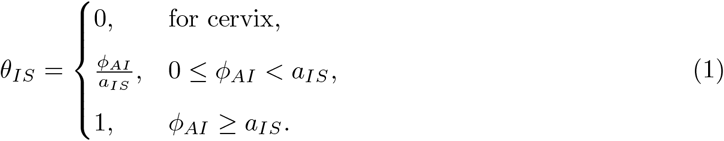

To achieve a physiological isthmus length that incorporated sufficient elements for a smooth transition of the material parameters, *a*_*IS*_ = 0.06 was used, resulting in an isthmus length of approximately 9 mm (Fig. 2a, bottom-right).

Additionally, we decomposed the combined uterocervical geometry into various segments along the anterior-posterior direction using the scalar field *ϕ*_*AP*_ to facilitate a quantitative comparison of the model-predicted deformation and strains (Fig. 2b). These included antero-medial (AM, *ϕ*_*AP*_ ∈ [0.59, 1.0]), antero-lateral (AL, *ϕ*_*AP*_ ∈ [0.51, 0.59)), postero-lateral (PL, *ϕ*_*AP*_ ∈ [0.47, 0.51)), and postero-medial (PM, (*ϕ*_*AP*_ ∈ [0.0, 0.47)) segments for the uterus, in addition to the cervix (Cx) as a separate segment (Fig. 2b).

#### 2.1.4 Fiber Generation

The uterus and cervix, like many other biological soft tissues, are fibrous [55, 56]. Fibers are organized protein networks, commonly composed of components such as collagen for tensile strength, elastin for tissue elasticity, and muscle fibers that define the direction of contraction. Fiber directionality is critical in characterizing the structure-function relationship of biological tissues. Although the evolution of fiber structures in the uterus and cervix throughout pregnancy is not fully understood, some information can be gathered from imaging methods such as optical coherence tomography (OCT) imaging [46, 57]. In the nonpregnant uterus, fibers are highly dispersed, contributing both circumferentially and longitudinally, while the nonpregnant cervix has predominantly circumferential fibers (more detailed spatial variation patterns described in Section 2.3), with dispersion that varies spatially. Alternatively, it was found that throughout pregnancy, the pregnant uterus transitions to predominantly longitudinal fibers, reinforcing the primary direction of uterine stretch, whereas the cervix maintains a predominantly circumferential (acting as a ratchet to control cervical dilation), but increasingly dispersed fiber structure [46, 57, 58].

For modeling purposes, a method to create smoothly varying fibers across the entire uterine tissue is essential to facilitate a smooth transition from the circumferentially aligned fibers in the cervix to the longitudinally aligned fibers in the uterus; however, such techniques have yet to be implemented. In cardiac mechanics modeling, rule-based approaches employing Laplace equations with suitably chosen Dirichlet boundary conditions are widely adopted to generate myocardial fibers [50, 51, 59–62].

Such approaches were effectively applied to simulate myocardial mechanics in the left and right ventricles, the left atrium, and even four-chamber whole heart models, yielding physiologically relevant cardiac motion and function [38, 43, 51, 60, 63–65].

We adopted the Laplace-Dirichlet rule-based (LDRB) algorithm to create spatially varying fiber directions that smoothly transition between the uterus and cervix throughout the isthmus region (Fig. 3). To do so, we used the previously introduced Laplace solutions (Section 2.1.2, Fig. 2a), *ϕ*_*TM*_ and *ϕ*_*AB*_, and the corresponding gradient fields to create a local orthonormal coordinate system at each mesh node as,

**Fig. 3.**
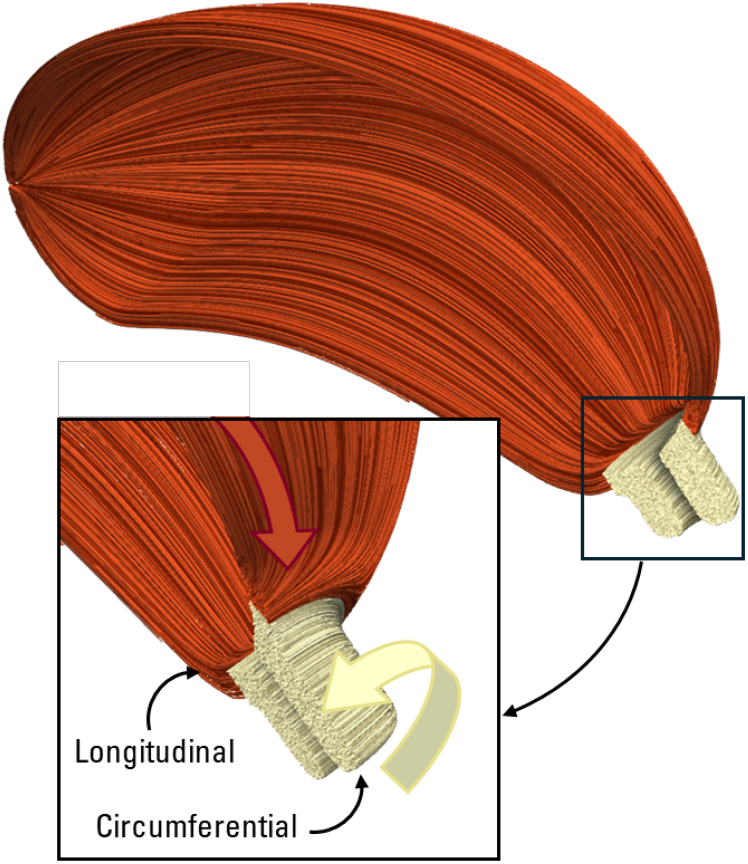
Rule-based uterine fiber structure. Rules are applied to the Laplace-Dirichlet solutions (*ϕ*_*TM*_, *ϕ*_*AB*_) to define fiber directions in the uterus and cervix. A convention is developed to allow a smooth transition from the longitudinal fibers in the uterus to the circumferential fibers in the cervix (Eqs. (2-3)). Please refer to Fig. 2a for the definitions of *ϕ*_*TM*_ and *ϕ*_*AB*_.

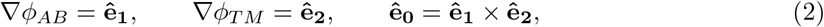

where **ê**_**0**_, **ê**_**1**_, **ê**_**2**_ are the local circumferential, longitudinal, and transmural coordinate directions, respectively. These basis vectors were then used to prescribe unique, yet smoothly transitioning fiber directions (**f**_**0**_) across the reference (i.e., unloaded) tissue domain as,

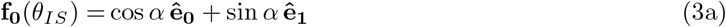

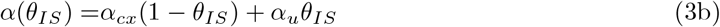

where *α*_*cx*_, *α*_*u*_ are constants and *θ*_*IS*_ is given in Eq. (1) (Fig. 2a, bottom-right). In this work, we set *α*_*cx*_ = 0 and *α*_*u*_ = *π/*2 radians to capture the circumferential and longitudinal fiber directions in the cervix and uterus, respectively (Fig. 3).

#### 2.1.5 Material Characterization

To model the uterine mechanical behavior, we adopted a modified form of the anisotropic, hyperelastic Gasser-Ogden-Holzapfel (GOH) constitutive model [45]. Although the GOH model was originally developed for two distinct fiber families, typically found in vasculature, here a single fiber family was sufficient to capture the experimentally characterized tissue mechanical response [46, 57, 66, 67]. The total strain energy density function (*ψ*) is then represented in its decoupled form, as,

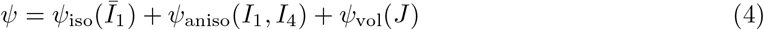

where Ψ_iso_, Ψ_aniso_ together represent the isochoric (volume-preserving) contribution, while *ψ*_vol_ contributes to changes in volume (dilation). Further, within the isochoric contribution, the isotropic component (Ψ_iso_) is a function of the isochoric invariant 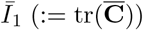, where as the anisotropic strain energy component (Ψ_aniso_) uses full invariants (*I*_1_ := tr(**C**), *I*_4_ := **f**_0_ **Cf**_0_) to correctly capture the volumetric-anisotropic deformations [68], with 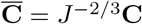 and **f**_0_ is the fiber direction in the reference configuration (Eq. (2)). Here, following the principles and definitions from nonlinear continuum mechanics [69], **C** := **F**^*T*^ **F** is the usual right Cauchy-Green deformation gradient tensor and the Jacobian, *J* := det(**F**) is the determinant of the deformation gradient tensor 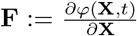. Further, φ : **Ω**_**X**_ → **Ω**_**x**_ is the deformation map between the material in its reference unloaded configuration **Ω**_**X**_ and the current deformed configuration **Ω**_**x**_, such that **x** = φ (**X**, *t*), where the material originally at **X** in the reference configuration is deformed to position **x** at time *t*. Lastly, **u** := **x**(**X**, *t*) − **X** is the displacement of a material particle. The specific forms of strain energy densities adopted in this work are given by,

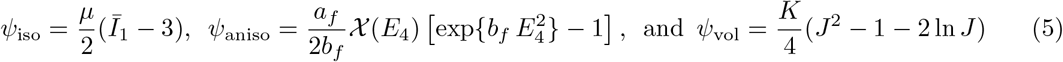

where *E*_4_ = *κI*_1_ + (1 3*κ*) − *I*_4_ − 1, with *κ* being the fiber dispersion parameter [45, 68]. In the above formulation,

i. The tissue ground substance was modeled using the isotropic neo-Hookean constitutive model (*ψ*_iso_ in Eq. (5)), parameterized by the shear modulus, 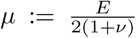, where *E* is the elastic modulus and *ν* is the Poisson ratio.
ii. The dispersive fibers were characterized using an exponential strain energy function (*ψ*_aniso_ in Eq. (5)), parameterized by fiber stiffness *a*_*f*_ and an exponential stiffening rate coefficient *b*_*f*_. While the original GOH model is formulated for collagen fibers under tension only that buckle under compressive loads [45], here, we allow a smooth transition between fiber distension and compression using 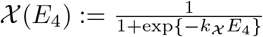, thus avoiding any numerical instabilities under small compressive strains [59, 60]. The parameter *k*_*X*_ controls the degree of smoothness, and is set to be *k*_*X*_ = 100 in this work.
iii. Finally, the volumetric deformations were penalized using the Simo-Taylor (ST91) constitutive model [70] (*ψ*_vol_ in Eq. (5)) parameterized using the material bulk modulus, 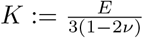.

#### 2.1.6 Material Parameter Tuning

The constitutive model parameters identified in the previous section for the uterus and the cervix were either derived from previous modeling approaches or optimized to match experimental data. Parameters *E, κ, a*_*f*_, and *b*_*f*_ for each domain were optimized to match sample-averaged force-stretch curves from experimental data of pregnant uterine and cervical tissues in tension and compression [66, 67].

The elastic modulus, *E*, was first estimated to match experimental data during compression, as the anisotropic terms do not contribute to the strain energy adopted here (Eqs. (4–5)). Therefore, to approximate the value of *E* required to capture the experimental spherical indentation test data, we adopted the Hertzian spherical contact mechanics model with the Dimitriadis correction for finite thickness [71]. The force-indentation depth relationship is given by,

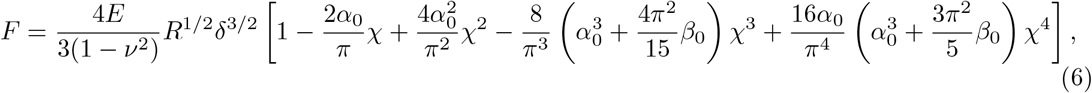

where 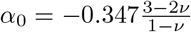, 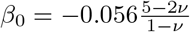, and 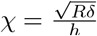. *F* is the force applied to the spherical indenter, *δ h* is the tissue thickness. is the indentation depth, *R* is the spherical indenter radius, and *h* values were chosen as the average of the experimental sample thicknesses (4.7 and 2.4 mm for uterus and cervix, respectively), and *R* values were chosen based on the spherical indenter used experimentally (3.0 and 1.25 mm for uterus and cervix, respectively) [66, 67]. Strain values were calculated as the ratio of indentation depth to full tissue thickness. The *E* values for the uterus and cervix were optimized using the SciPy curve-fit function, minimizing the sum of squared residuals between the model fit and experimental data.

Fiber parameters *k, a*_*f*_, and *b*_*f*_ were subsequently optimized to refine the model fit and match the experimental data under tension for the uterus and cervix. The constitutive model (Eq. (5)) was analytically solved for a uniaxial tensile test with fibers aligned in the loading direction and traction-free lateral boundaries. Simulated tissue cross-sectional areas were based on average dimensions of the experimental samples [66, 67]. Again, the SciPy curve-fit function was used to optimize the parameters to match the experimental data. The Poisson ratio, *ν*, was selected based on previous modeling approaches and was found to have minimal influence on the overall model fit [19].

Using the optimized parameters (Table 1) resulted in a strong agreement with the experimental data for both uterus and cervix during tension and compression (Fig. 4b), with the standard errors for the uterus and cervix being 0.0019 N and 0.0007 N, respectively. To capture the variability in the experimentally measured force-stretch curves, however, we varied the material parameters around these fit baseline values as part of the sensitivity analysis (Section 2.3). These baseline material parameters are subsequently used to model the passive mechanics of the image-based uterus model to optimize boundary conditions, as discussed next.

**Table 1.**
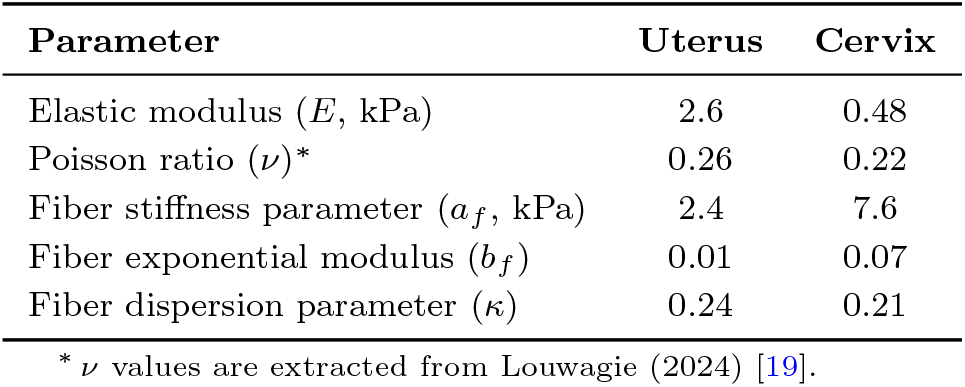
Iteratively optimized material parameters to fit *in vitro* mechanical testing data of Fang et al. [67] for the uterus and Shi et al. [66] for the cervix.

| Parameter | Uterus | Cervix |
| --- | --- | --- |
| Elastic modulus ($E$, kPa) | 2.6 | 0.48 |
| Poisson ratio ($\nu$)* | 0.26 | 0.22 |
| Fiber stiffness parameter ($a_f$, kPa) | 2.4 | 7.6 |
| Fiber exponential modulus ($b_f$) | 0.01 | 0.07 |
| Fiber dispersion parameter ($\kappa$) | 0.24 | 0.21 |

\* $\nu$ values are extracted from Louwagie (2024) [19].

**Fig. 4.**
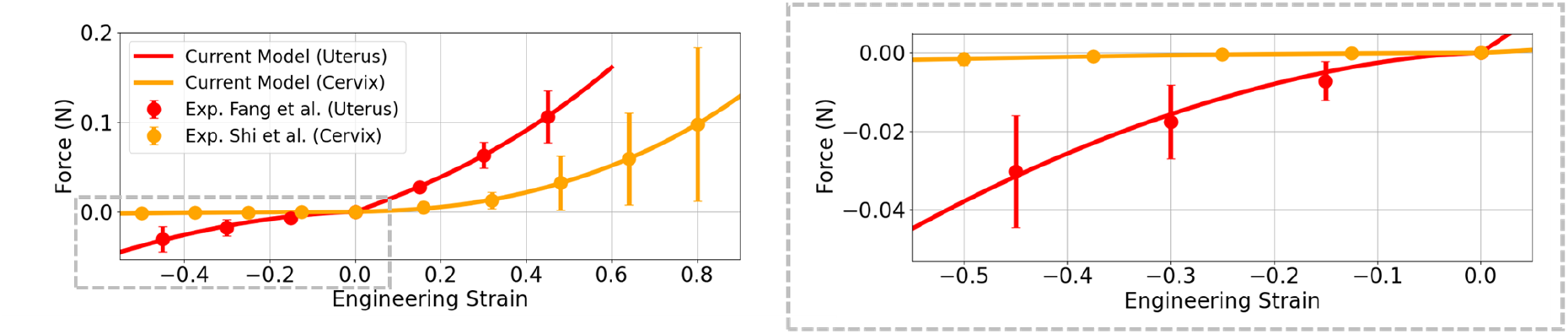
Material parameter estimation using *in vitro* mechanical test data. (left) Comparison of the current model fit against experimental data extracted from Fang et al. [67] for the uterus and from Shi et al. [66] for the cervix during pregnancy. (right) A zoomed-in view of the model fit during compression.

#### 2.1.7 Boundary Condition Development

The current state-of-the-art approach for modeling uterine passive mechanics is configured in FEBio^2^ [72], developed by Louwagie (2024) [19], which has been refined for improved fidelity from prior work [47]. Although it was originally developed for second-trimester modeling, it has been adopted by other groups to study active [21] and passive [26, 28] uterine mechanics at term. This approach relies on modeling contact with adjacent tissue components (abdomen and fetal membrane) to constrain the motion of the uterus and cervix under physiological loads (Fig. 5a, left). While this approach is powerful in capturing passive mechanics with reasonable biofidelity, incorporating these additional tissue components and contact requirements limits the model’s robustness and efficiency, which is necessary for the long-term goals of this work, including longitudinal growth and remodeling and coupled excitation-contraction studies. Therefore, we sought to employ this approach to create our verification dataset, aiming to achieve similar deformations under passive loading by introducing a more computationally efficient treatment of boundary conditions for the uterus and cervix (Fig. 5a, right).

**Fig. 5.**
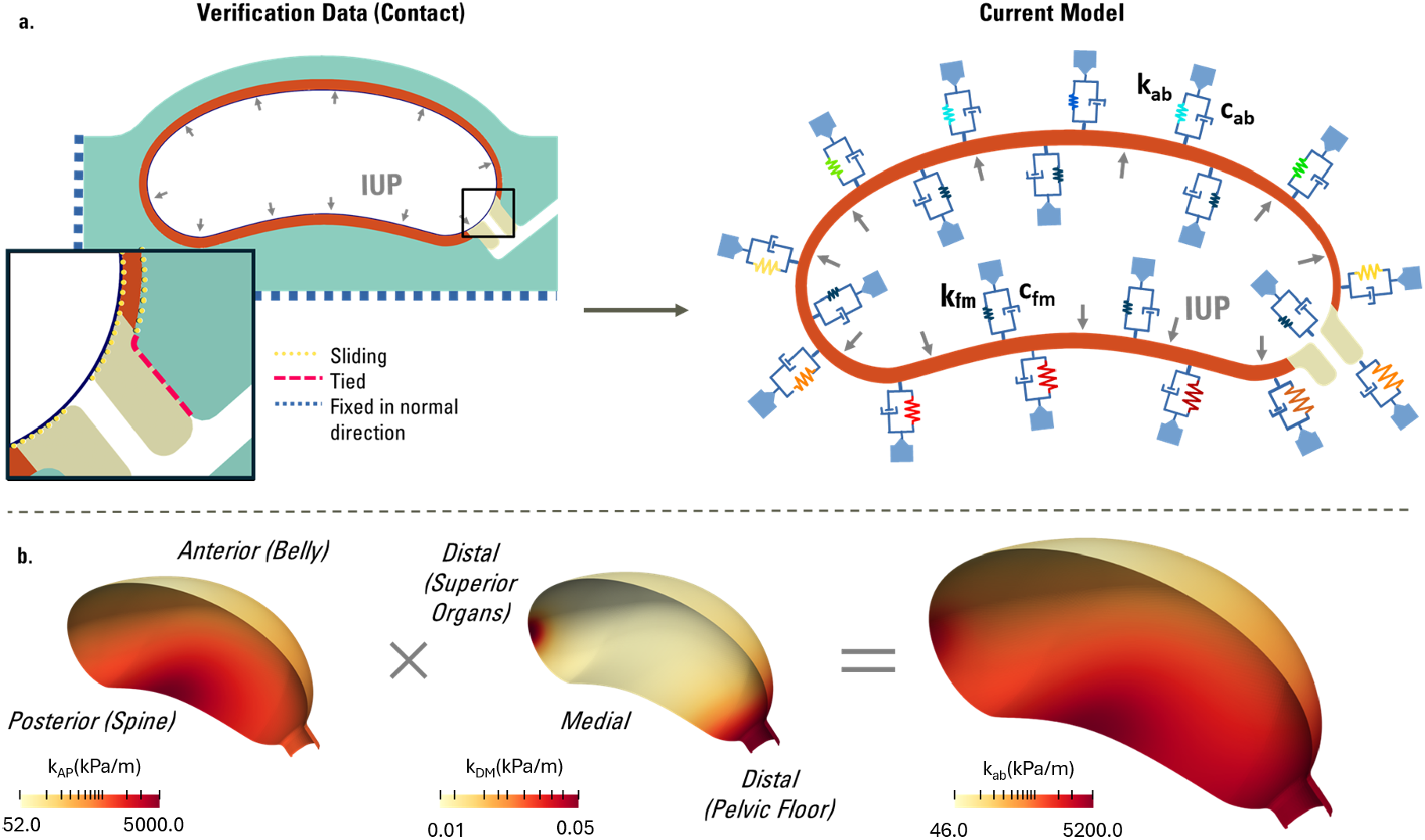
Boundary condition framework development to simulate uterine passive mechanics. (a) (left) Schematic of boundary condition setup along the mid-sagittal plane view used in a well-established FEBio model of uterine mechanics that involves different types of contact surfaces. Tied contacts were used on the uterus-cervix and cervix-abdomen interface. The uterus was allowed to slide within the abdomen. The fetal membrane was tied for approximately 70 % of the superior uterus-membrane interface and allowed to slide through the remaining 30 % on the inferior side, a combination found optimal in Louwagie (2024) [19]. The membrane was allowed to slide along the top surface of the cervix. The posterior and superior surfaces of the abdomen were fixed in their normal directions, and symmetry boundary conditions were applied along the mid-sagittal face. (right) Schematic of the current model with Robin boundary conditions involving linear springs along the outer and inner surfaces of the uterus and cervix with spatially varying coefficients. An intra-uterine pressure (IUP) was applied to the internal surface of the fetal membrane for the contact-based FEBio model (left) and the uterine inner surface for the current model (right). (b) (left) Spatially varying stiffness coefficients along the (left) anterior-posterior (*k*_*AP*_) and (center) distal-medial (*k*_*DM*_) directions. Anatomical directions and the surrounding organs represented by the applied boundary conditions are labeled. (right) The overall stiffness coefficient *k*_*ab*_, computed as the product of *k*_*AP*_ and *k*_*DM*_, is representative of the effective stiffness acting on the uterus due to the abdomen and other neighboring tissues.

To create this verification dataset in FEBio, we implemented identical patient geometry, mesh, fibers, and material properties (using the HGO Unconstrained constitutive model^3^) for the uterus and cervix. The only difference with the current approach, therefore, is in the boundary conditions. The abdomen and fetal membrane were added following the modeling framework provided in Louwagie (2024) [19]. The abdomen is made up of a neo-Hookean material, and the fetal membrane is made up of a neo-Hookean ground substance with transversely isotropic, continuously distributed embedded fibers. Contact conditions were chosen based on previous approaches, implementing physiologically relevant combinations of tied-elastic and sliding-elastic contacts (Fig. 5a, left). A pressure of 1.7 kPa was applied to the internal surface of the fetal membrane, chosen to match physiological intrauterine pressures (IUP) at 38 weeks of gestation [73]. All other FEBio solver settings are summarized in Table 3.

To improve model robustness, we then replaced the abdomen and fetal membrane components with computationally efficient Robin boundary conditions (Fig. 5a, right). These boundary conditions have been widely implemented in the cardiovascular mechanics field due to their ability to capture interactions with unmodeled tissue components by simulating springs and dashpots on the relevant model surfaces [36, 38, 41, 43, 44, 74]. Here, we employed this technique to capture the motion constraints imposed by surrounding organs and the fetal membrane (Fig. 5a, right). These conditions are mathematically expressed as,

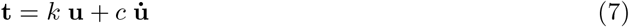

where **t** is the traction vector on the outer surface of the uterus and cervix, *k* is the spatially-varying omnidirectional stiffness value, and *c* is the damping coefficient. **u** and 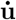 are the local material displacements and velocities, respectively, defined in Section 2.1.5.

The effective stiffness introduced by the abdomen varied spatially, depending on the thickness and boundary conditions of the abdomen configuration employed in the contact-based model. To capture this, we applied spatially varying Robin stiffness coefficients (*k*_*ab*_) to the outer surface of the uterus and cervix (Eq. (9), Fig. 5b). We defined *k*_*ab*_ as a product of two smooth nonlinear stiffness distributions along the anterior-posterior direction (*k*_*AP*_) and the distal-medial sector of the apico-basal direction (*k*_*DM*_). A constant, non-zero damping coefficient is applied along the abdominal interface for numerical stability (Table 2). In what follows, we will describe our procedure for obtaining the spring stiffness distributions.

**Table 2.** Tuned boundary condition parameters to match the deformation of the contact-based verification model.

| Parameter Type | Parameter | Value |
| --- | --- | --- |
| Mapping Parameters | $a_{DM}$ | -12 |
| | $a_{AP}$ | -22 |
| | $b_{DM}$ | -0.50 |
| | $b_{AP}$ | -0.38 |
| Stiffness Parameters | $k_D$ (kPa/m) | $5 \times 10^{-2}$ |
| | $k_M$ (kPa/m) | $1 \times 10^{-2}$ |
| | $k_A$ (kPa/m) | 52.0 |
| | $k_P$ (kPa/m) | $5 \times 10^3$ |
| | $k_{fm}$ (kPa/m) | 6.5 |
| | $c_{ab}$ (kPa·s/m) | 35.0 |
| | $c_{fm}$ (kPa·s/m) | 0 |

**Table 3.** Summary of the solver features and key parameters for the current model and verification data.

| Solver Feature/Setting | Current Model (*svMultPhys*<sup>4</sup>) | Verification Dataset (FEBio<sup>7</sup>) |
| --- | --- | --- |
| Parallelism | Domain Decomposition<br>(MPI-based) | Shared Memory Parallelism<br>(OpenMP) |
| Analysis | dynamic | quasi-static |
| Time integration | generalized-α | – |
| Spectral radius of<br>infinite timestep ($\rho_\infty$) | 0.5 | – |
| Time stepping | uniform ($\Delta t = 0.01$) | auto stepper* ($\Delta t \in [10^{-6}, 0.1]$) |
| Nonlinear solver | full Newton | quasi-Newton (BFGS) |
| Relative tolerance | $10^{-6}$ | $10^{-3}$ |
| Max Newton iterations | 10 | ∞ |
| Linear solver | GMRES (PETSc<sup>6</sup>) | direct (Pardiso) |
| Preconditioner | geometric AMG<sup>5</sup> (PETSc<sup>6</sup>) | – |

\*For quasi-static modeling, FEBio uses time-stepping for load increments only, rather than time integration.
<sup>4</sup>https://github.com/SimVascular/svFSI
<sup>5</sup>https://petsc.org/release/manualpages/PC/PCGAMG/
<sup>6</sup>Portable, Extensible Toolkit for Scientific Computing (PETSc), version 3.18.5, https://petsc.org/release/
<sup>7</sup>FEBio version 3.5.0, https://febio.org/

The Laplace solutions *ϕ*_*AB*_ and *ϕ*_*AP*_ (defined in Fig. 2a) along the outer surface of the uterus and cervix of were mapped to create exponential scale factors, *θ*_*AP*_ and *θ*_*DM*_, as,

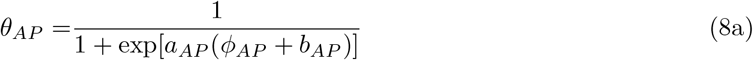

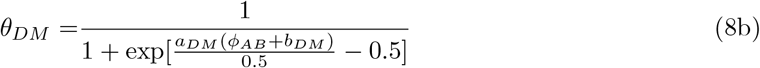

where *θ*_*AP*_ maps *ϕ*_*AP*_ to an exponentially varying scaling factor from the posterior to anterior uterine wall (Eq. (8a)), and *θ*_*DM*_ maps *ϕ*_*AB*_ to an exponentially varying scaling value from the medial to distal regions of the uterus and cervix (Eq. (8b)). Parameters *a*_*AP*_ and *a*_*DM*_ control the steepness of the exponential function, while *b*_*AP*_ and *b*_*DM*_ control the lateral shift of the function. These scale factors were then used to obtain the nonlinear stiffness distributions,

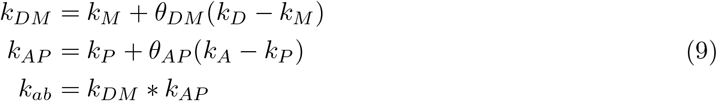

where *k*_*M*_, *k*_*D*_ are the stiffness bounds along the medial and distal directions, while *k*_*A*_, *k*_*P*_ are the bounds along the anterior-posterior direction.

Further, a uniform stiffness (*k*_*fm*_) was applied to the internal surface of the uterus and cervix to capture the constraint due to the fetal membrane. This stiffness value is relatively small and was chosen based on the relative stiffnesses of the anterior abdomen and fetal membrane, as determined by spherical inflation tests (Fig. A1). Here, two hollow spheres, resembling the idealized abdomen and fetal membrane, were created with the same inner diameters, matching the perpendicular distance from the inner surface of the anterior wall of the uterus at the anterior-most point of the uterus to the inner surface of the posterior wall of the uterus (*d*_*U*_ in Fig. A1). The wall thicknesses were chosen based on the thickness of the anterior abdominal wall at the peak of the belly (*t*_*ab*_ = 20 *mm*) and the fetal membrane (*t*_*fm*_ = 0.8 *mm*). The material properties used for these tissues in the FEBio model were applied to the respective spheres and inflated with an IUP of 1.7 kPa. Relative stiffnesses of the tissues were then computed based on respective average displacement through the tissue thickness after inflation. The ratio of fetal membrane to abdominal stiffness was found to be ~ 1/8; therefore, this value was used to estimate the scale of the fetal membrane stiffness from *k*_*A*_. The damping coefficient along the fetal membrane, *c*_*fm*_, was set to zero for modeling simplicity.

We then applied the same pressure load of 1.7 kPa to the internal surfaces of the uterus and cervix as in the FEBio model. Using the material parameters fit to match experimental data (Table 1) and finite element procedure described in Section 2.2 (Table 3), we manually tuned the Robin boundary condition mapping parameters (*a*_(·)_, *b*_(·)_ in Eq. (8)) and stiffness values (*k*_(·)_ in Eq. (9)) to match the deformation captured in the FEBio model. Errors are quantified in terms of the L2 norm of the differences in nodal displacements between the two approaches locally and globally and further verified by comparison of Lagrange strain (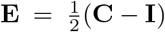) components (Table 5 in Section 3.1, Table A2 and Fig. A2 in Appendix). The combination of these tuned boundary condition parameters (Table 2) ultimately resulted in higher stiffness where the uterus is in contact with the pelvic floor, abdominal organs superior to the uterus (i.e., bowel, liver, stomach), and spine, but allowed greater distensibility along the anterior surface of the uterus (Fig. 5b).

### 2.2 Finite Element Solver

The simulations developed here were performed using an in-house multiphysics finite element solver, *svMultPhys*, adapted from the open-source finite element solver, *svFSI* ^4^ [75], which has been previously verified for cardiac mechanics applications [59, 76], and validated and employed for other cardiovascular biomechanics applications, including patient-specific multiscale cardiac mechanics modeling [38, 59, 60, 77], cardiac electrophysiology [78], myometrial excitation during pregnancy [14], blood flow in coronaries [79], developing ventricles [80, 81], fluid-structure interaction modeling in aortic dissection and aneurysms [39, 41, 82], and multiphysics applications [83], including growth and remodeling [84, 85]. The finite strain elastodynamics equations governing uterine passive mechanics were solved using the standard Galerkin approach with displacement-based formulation and continuous finite element basis [86]. Time integration was performed using the implicit, second-order accurate, generalized-*α* method. The resulting nonlinear system of equations was solved using the Newton-Raphson method, embedded within a multi-step predictor-corrector method [87]. Within each nonlinear iteration, the linearized system of equations is solved using the Generalized Minimum Residual (GMRES) iterative solver, preconditioned with the geometric algebraic multigrid method (GAMG^5^), invoked from within the PETSc library^6^. Table 3 summarizes key solver parameters and other features and settings employed in the current study and in creating the FEBio-based verification dataset.

### 2.3 Sensitivity Analysis

#### 2.3.1 Material Parameter Bounds

Although the material parameters were reasonably fit to match mechanical testing data (Section 2.1.6), experimental data suggested substantial variability in the mechanical response across the tissue samples (Fig. 4b). This raises two questions:

1. First, what are the most influential material parameters and their ranges that can reasonably capture the observed experimental behavior in both tension and compression?
2. Second, how much do these parameter variations impact the passive mechanical response of the three-dimensional parametric model of the uterus under physiological pressures?

To address these questions, we performed a sensitivity analysis by varying material parameters around the baseline values identified in Table 1. Our goal here was to determine parameter bounds that reasonably capture the observed spread in the experimental force-stretch relationship for both the uterus and the cervix, under both tension and compression. We then simulated the uterine passive mechanics by systematically varying these parameters within their bounds to evaluate changes in its local and global biomechanical behavior. Additionally, we assessed the biomechanical impact of the heterogeneity in the cervical fiber structure documented in previous studies [88, 89].

To determine the material parameter bounds, we focused on varying the elastic modulus of the ground substance, *E*, and fiber parameters (*a*_*f*_ and *κ*). *E* was varied to capture the experimental variability in compressive mechanical behavior of the uterus and cervix (Fig. 4b). The contributions of fibers were also varied through adjustments of the fiber stiffness, *a*_*f*_, and the fiber dispersion parameter, *κ*. For example, reducing the circumferential fiber contribution could be captured through a decrease in *a*_*f*_ or an increase in *κ* locally. For each of these parameter variations, we characterized the force-strain relationship for tension and compression using the analytical parameter-fitting protocol presented in Section 2.1.6; however, parameters were fit to the experimental data points that define the standard deviation bounds, rather than to the average.

Table 4 reports the parameter bounds that reasonably captured the variance in the force-strain response for both uterus and cervix, across tension and compression (Fig. 6a). In particular, varying fiber parameters captured the variability in tensile characteristics, while modulating ground substance *E* captured the spread mainly during compression (Fig. 6a). Due to the larger fiber contribution in the cervix compared to the uterus, changes in *E* resulted in smaller variations in the cervical tensile characteristics relative to the uterus.

**Table 4.** Material parameter bounds to capture the experimental variability in the force-strain response (Figures 4b, 6a). (%) indicates relative change from the baseline value reported in Table 1.

| Parameter | Uterus |  |  | Cervix |  |  |
| --- | --- | --- | --- | --- | --- | --- |
|  | Baseline | Upper Bound | Lower Bound | Baseline | Upper Bound | Lower Bound |
| $E$, kPa | 2.6 | 3.9 (+49 %) | 1.3 (-49 %) | 0.48 | 0.92 (+91 %) | 0.23* (-53 %) |
| $a_f$, kPa | 2.4 | 3.6 (+45 %) | 1.4 (-42 %) | 7.6 | 15.0* (+97 %) | 0.70 (-91 %) |
| $\kappa$ | 0.235 | 0.28 (+19 %) | 0.20 (-15 %) | 0.214 | 0.33<sup>†</sup> (+54 %) | 0.15 (-30 %) |

\* Limited by model convergence due to excessive $a_f/E$ ratio
<sup>†</sup> Limited to the upper bound of dispersion, i.e., $\kappa = 1/3$ , representing a fully dispersed fiber structure.

**Fig. 6.**
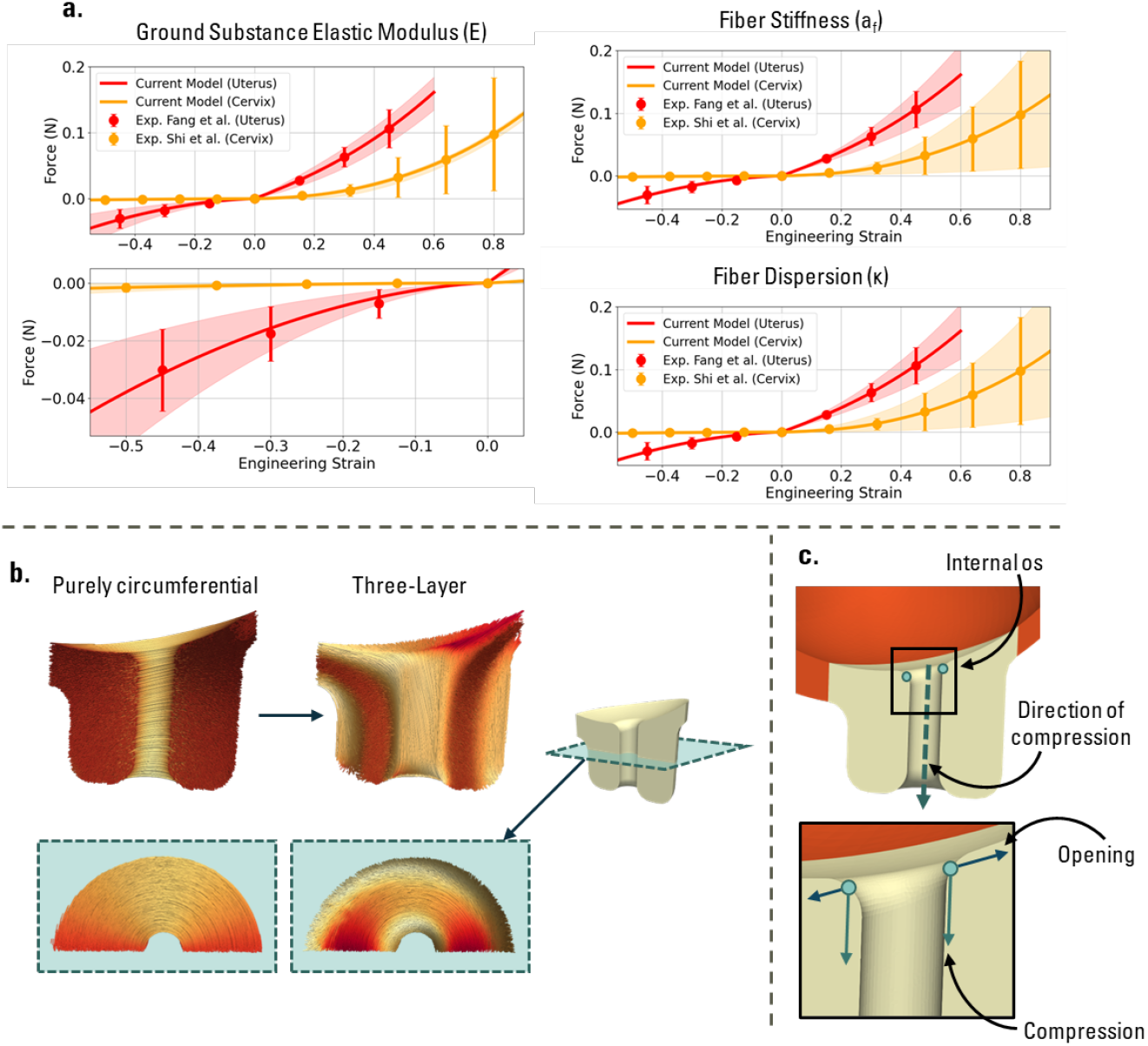
Sensitivity analysis parameterization. (a) Simulated sensitivities of the force-stain relationship to variations in tissue material parameters, including (left) ground substance elastic modulus, *E*, (center) fiber stiffness parameter, *a*_*f*_, and (right) fiber dispersion parameter, *κ*, compared against variability in the mechanical testing data of Shi et al. [66] for the cervix and Fang et al. [67] for the uterus. (b) Changing the cervical fiber structure from purely circumferential to a three-layered structure. (c) Schematic of internal ostium (os) for calculating cervical opening and compression.

We also sought to verify that the parameter ranges identified as spanning the variability in the experimental force-strain data were realistic. To this end, we simulated tension and compression tests for each sample’s force–strain data, extracted from their original studies [66, 67], rather than using the sample-averaged profile (Section 2.1.6). We found that all estimated sample-specific parameter values were within the bounds determined for sensitivity analysis (Table 4), confirming their physiological relevance.

#### 2.3.2 Fiber Architecture

Further, the cervix fiber directionality and dispersion were mostly quantified along the circumferential direction, motivating the use of a preferential circumferential fiber architecture within our model (Section 2.1.4) [57]. However, anecdotal observations and other imaging studies have described a three-layered fiber structure in the cervix, where longitudinal fibers make up the outer and inner surfaces of the cervix (parallel to the endocervical canal), with circumferential fibers in between [58, 88, 89]. Therefore, to capture this pattern (Fig. 6b), we used the transmural Laplace solution, Φ_*tm*_ (Fig. 2a, top-left), to define a sinusoidal scalar field 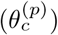 cervical tissue as,

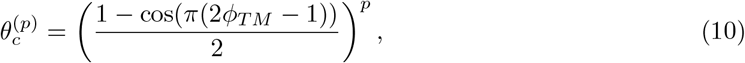

where, *p* controls the proportion of circumferential and longitudinal fibers. This scalar field was then used to prescribe the fiber angles as,

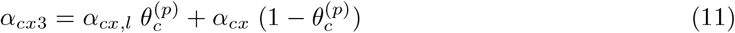

where, *α*_*cx,l*_ = *π/*2 radians to capture the longitudinal fibers on the external and internal surfaces of the cervix, while *α*_*cx*_ was set to 0 (Eq. (3b)). These fiber angles were then mapped to nodal fiber directions using orthonormal basis vectors defined in Eqs. (2-3). In this work, we chose *p* = 1 based on published diffusion tensor MRI data of the cervical fiber architecture [89] (Fig. 6b).

#### 2.3.3 Analysis Metrics

To quantify the mechanical characteristics of the passive uterus in response to variations in material parameters (Table 4) and fiber structure (Fig. 6b), we measured percent changes in Green-Lagrange strain magnitude (calculated as Frobenius norm of the Green-Lagrange strain tensor), cervical opening, and cervical compression. Changes in strain concentrations at the internal os (internal ostium or the internal opening of the cervix, IO, Fig. 6c) were quantified as the average percent change in strain magnitude across elements encircling the internal os. The percent change in cervical opening was measured as the increase in distance between two points on the mid-sagittal plane of the internal os relative to the baseline model (Fig. 6c). The degree of compressibility of the cervix was quantified by the percent change of the displacement of the internal os in the direction of the undeformed cervical canal relative to the baseline configuration (Fig. 6c). Lastly, changes in the Lagrange strain magnitude throughout the uterus were measured as percent differences in the averaged strain relative to the baseline model.

## 3 Results

### 3.1 Model Configuration

A summary of all the parameters, including tuned material parameters fit to experimental data (Table 1), optimized boundary condition parameters (Table 2), and solver settings (Table 3), used to create the baseline model of the uterine passive mechanics, is provided in the appendix (Table A1). Using this parameter set, the nodal differences in uterine displacements between the current model and contact-based model under physiological loading averaged 3.4 ± 1.8 mm across both the uterus and the cervix, with a maximum difference of 7.4 mm (Fig. 7). Separately, the cervix showed much closer agreement with the contact-based model compared to the uterus (1.0 ± 0.2 mm for cervix vs. 3.7 ± 1.7 mm for the uterus, Table 5). It should be noted that, despite a close geometric agreement between the two deformed configurations, some nodal displacement differences persisted due to sliding motion. This effect is apparent in the posterior region, where the geometries aligned well, yet differences in in-plane sliding of the uterus ultimately led to larger differences in nodal displacements (Fig. 7b, left). Region-wise partitioning of the uterus (Fig. 2b) revealed the most substantial differences between the two approaches at the lateral wall, across both the anterior (AL) and posterior (PL) regions (Table 5). Spatial variations in strain components demonstrated reasonable agreement between the two modeling approaches, with the region-averaged nodal differences between strain components all below 0.065 (Fig. A2, Table A2).

**Fig. 7.**
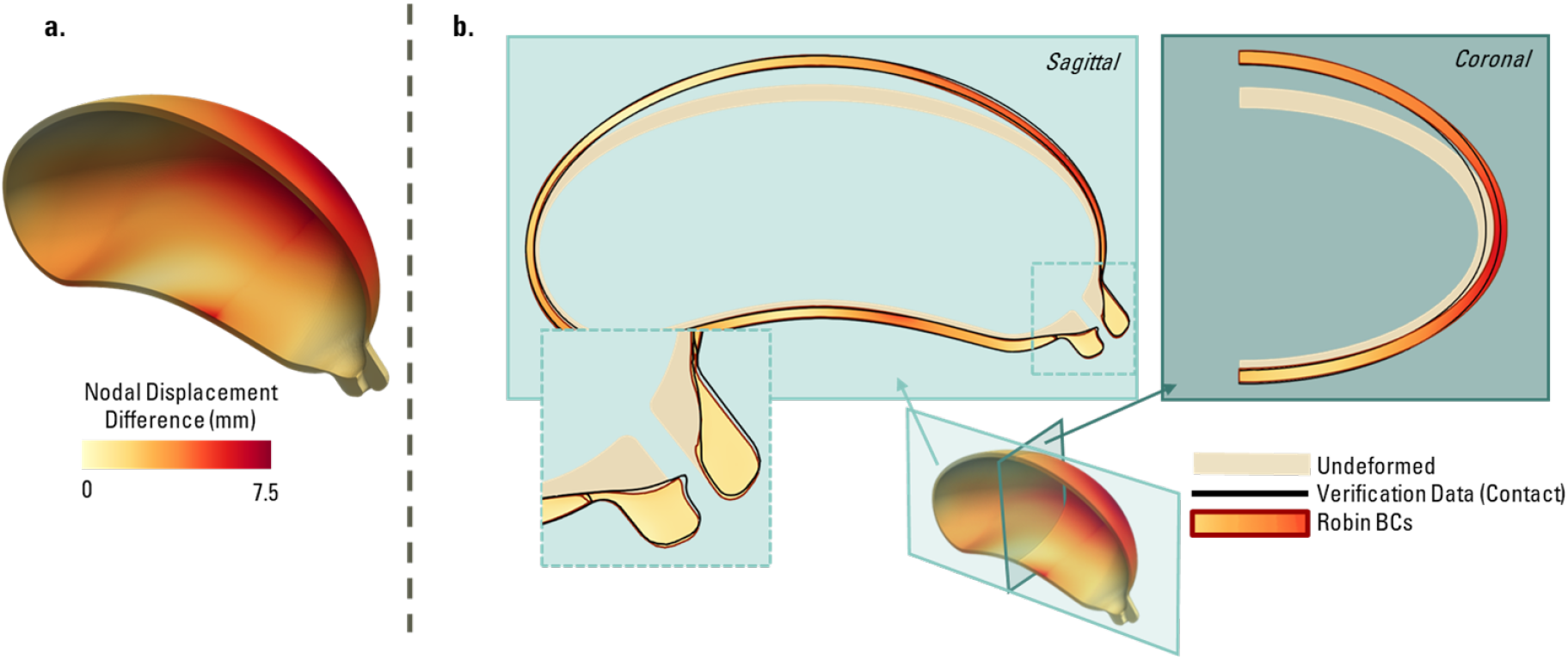
Model verification. Comparison of the mechanical deformation predicted by the current model employing Robin boundary conditions against a previously established workflow using contact with surrounding organs, employed here as a verification dataset (Fig. 5a, left). (a) A sectional view of the uterus showing a map of the nodal differences in displacements. (b) Views of the deformed model from each modeling approach along the mid-sagittal (left, lighter green background) and mid-coronal (right, darker green background) planes.

**Table 5.** L2 norm of the nodal differences in displacements (Err_**u**_) between the two modeling approaches. Values reported as regional mean *±* standard deviation.

| Tissue Domain | Region* | $\text{Err}_u$ (mm) |
| --- | --- | --- |
| Uterus | Overall | $3.7 \pm 1.7$ |
| | Antero-Medial (AM) | $3.1 \pm 1.4$ |
| | Antero-Lateral (AL) | $4.3 \pm 2.1$ |
| | Postero-Medial (PM) | $3.0 \pm 1.0$ |
| | Postero-Lateral (PL) | $4.3 \pm 1.4$ |
| Cervix (Cx) | | $1.0 \pm 0.2$ |
| Combined Model | | $3.4 \pm 1.8$ |

\*Please refer Fig. 2b for more details on uterus partitions.

The current approach, however, resulted in substantial savings in computational time, by nearly 90 %. The total simulation time using the current modeling approach with Robin boundary conditions was 1940 seconds, while the FEBio model, which included contact with surrounding tissues, took 18,422 seconds. The FEBio model employs an auto-timestepper, enabling the simulation to be solved in 22 timesteps. In contrast, the current solver, *svMultPhys*, remains limited to uniform time stepping, requiring 100 time steps for this simulation.

### 3.2 Sensitivity Analysis

Varying the elastic modulus of the ground substance and fiber properties resulted in substantial changes in the mechanical behavior of the uterus and cervix (Table 6 and Fig. 8). The uterine tensile strains were found to be less sensitive to fiber properties, but showed a strong inverse relationship with the tissue elastic modulus (Table 6). Compared to the uterus, however, the cervical mechanics were found to be sensitive to all the material changes considered in this study (Table 6). A 91% increase in *E* in the cervix resulted in a decrease in strain concentration in the internal os (−9.8 %), cervical opening (−9.1 %), and cervical compression (−19.2%). A proportional increase in these mechanical indices was also found when reducing *E*. Alternatively, a disproportionate effect was found when increasing or decreasing *a*_*f*_. Lowering *a*_*f*_ resulted in a substantial increase in cervical mechanical measures compared to smaller reductions in each of these measures when *a*_*f*_ is increased proportionally. A similar disproportionate effect was found when modulating *κ*, but to a lesser extent. *κ* has a strong effect on the internal os strain (−15.7 % to +74.7 %) and cervical opening (−29.0 % to +88.9 %), but has a weak impact on cervical compression (−0.9 % to +4.9 %). Lastly, changing the fiber architecture to the three-layered model substantially increases internal os strain and cervical opening, but has a reduced effect on cervical compression (Table 6).

**Table 6.** Characterizing changes in passive mechanical deformation of the uterus and cervix due to variations in material parameters (Table 4) and cervical fiber structure (Fig. 6b). All values are reported as percentage difference from the baseline model.

| Mechanical Property<sup>§</sup> | Uterine Strain (%) | Internal Os Strain (%) | Cervical Opening (%) | Cervical Compression (%) |
| --- | --- | --- | --- | --- |
| $\uparrow E$ (+49%, +91%) | -17.9 | -9.8 | -9.1 | -19.2 |
| $\downarrow E$ (-49%, -53%*) | 27.9 | 5.6 | 5.85 | 13.6 |
| $\uparrow a_f$ (+45%, +97%*) | 0.0 | -6.7 | -13.5 | -2.9 |
| $\downarrow a_f$ (-42%, -91%) | 0.0 | 68.0 | 91.9 | 19.9 |
| $\uparrow \kappa$ (+19%, +54%<sup>†</sup>) | 0.0 | 74.7 | 88.9 | -0.9 |
| $\downarrow \kappa$ (-15%, -30%) | 0.0 | -15.7 | -29.0 | 4.9 |
| Three-Layer Model | 0.0 | 100.3 | 121.0 | -7.5 |

<sup>§</sup> Parameters are either simultaneously set to their maxima or minima for both the uterus and cervix. The first element of the ordered pair is for the uterus, and the second element is for the cervix.
\*Bounds limited by model convergence due to excessive $a_f/E$ ratio.
<sup>†</sup>Bound limited by upper limit of $\kappa = 1/3$ , representing fully dispersed cervical fibers.

**Fig. 8.**
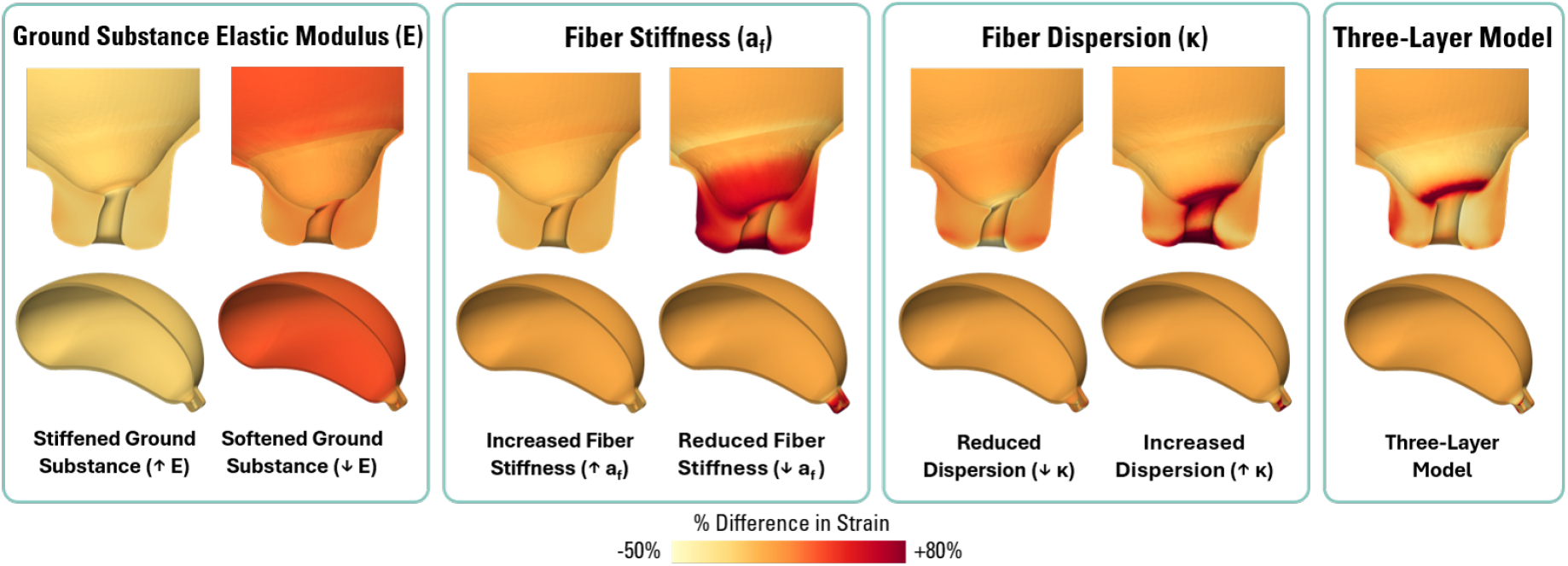
Sensitivities to material parameters and fiber structure. Relative local differences in strain with respect to the baseline model across the uterus and cervix in response to changes in the material properties, including (left to right) ground substance elastic modulus (*E*), fiber stiffness (*a*_*f*_), fiber dispersion (*κ*), and fiber structure (Table 4, Fig. 6). All simulations were performed in the current Robin boundary condition configuration.

These effects can also be qualitatively observed in Fig. 8. For changes in *E* and *a*_*f*_, both changes in strain concentrations at the internal os and overall cervix strains are visible (yet small in the case of increased *a*_*f*_ due to disproportionate effect discussed earlier). However, in the case of changes in *κ* and the three-layer model, strain concentrations at the internal os are apparent, but changes in global cervix strains are minimal.

## 4 Discussion

Uterine mechanics synthesizes a complex system of electro-chemo-mechanical pathways that tightly regulate dramatic tissue evolution throughout pregnancy [9]. These processes span large time scales, with tissue growth and remodeling progressing over months, contraction occurring a few seconds or minutes, and labor lasting hours. Further, many length scales are also relevant, ranging from cell-level action potential propagation to tissue-level growth to organ-level interactions [9]. Such separations of time and length scales introduce significant computational challenges. Therefore, we sought to develop a passive mechanics modeling workflow for a pregnant uterus, built upon a recently developed parametric model creation pipeline (Section 2.1.1) [19, 22] that prioritizes model robustness and efficiency for future longitudinal and active mechanics studies.

In this work, we introduced a novel workflow that cross-fertilizes cardiac mechanics modeling approaches with uterine passive mechanics (Section 2.1). For the first time, we adapted Laplace-Dirichlet rule-based approaches for creating fibers across the myocardium to the uterus and cervix, guided by OCT imaging and other anecdotal observations (Section 2.1.4, Fig. 3). To model the mechanical response of these fibers, we have adapted the hyperelastic GOH constitutive model, developed for vascular mechanics, with an exponential dependence on fiber strains while accounting for fiber dispersion (Section 2.1.5, Eq. (5)). Similar to the GOH model, the ground substance is assumed to be isotropic and have a linear dependence on the strain, represented using the neo-Hookean model (Section 2.1.5, Eq. (5)). Further, through the implementation of Robin boundary conditions informed by Laplace-Dirichlet solutions (Section 2.1.7), we were able to capture the displacements achieved in a contact-based model within reasonable agreement (Table 5, Fig. 7). With this approach, the computational expense of contact and additional adjacent tissue components is eliminated, ultimately reducing the computational time of the passive mechanics simulations to about 1*/*10^*th*^ of the previously established model (Section 3.1), while acknowledging major differences between the two solvers and solution procedures (Table 3).

Through this model, we were also able to explore the changes in the passive mechanics of the tissue in response to altered mechanical properties within physiological ranges (Sections 2.3, 3.2). Cervical softness has been found to be a predictor of preterm birth both through clinical elastography and *in vivo* stiffness assessments using aspiration-based devices [90–92] and shear wave elastography [93–95]. Cervical softness was captured in this model by reducing fiber stiffness (*a*_*f*_) and the elastic modulus of the ground substance *E*. Lowering *a*_*f*_ demonstrated substantial changes in the cervical deformation, both through enabling increased opening of the cervix and increased compression (Table 6, Fig. 8). This may occur physiologically due to weakened collagen crosslinking, higher presence of immature collagen, or increased collagen degradation [58]. Similar changes in the deformation of the cervix were found when varying *E*, representative of altered proteoglycan, glycoproteins, and glycosaminoglycan composition of the tissues [58]. The material is characterized such that only the ground matrix properties (*E*) govern tissue compression, while both the ground matrix and fiber properties impact the tissue’s mechanical response during tension. However, in the sensitivity analysis, both fiber stiffness and elastic modulus facilitate the degree of cervical compression. Such findings highlight the complex loading state of the cervix. When the cervix compresses, the generated strain field around the internal os facilitates the cervical opening. Therefore, circumferential fibers are recruited in tension, modulating the overall deformation of the tissue. The increased cervical strain upon softening observed in this model can be placed in the context of inflammatory pathways of cervical ripening. Experimental studies have suggested that mechanical signaling may contribute to cervical ripening, and, in turn, affect labor timing [12]. Premature or insufficient cervical ripening may contribute to preterm or post-term births, respectively [7, 12]. As such, this work reinforces the potential mechanical role of pathological cervical softening in the increased risk of labor complications.

Further, this work has also introduced an investigation of cervical fiber organization (Fig. 6b) through both fiber dispersion and overall fiber architecture (three-layer model). From our results (third and fourth columns of Fig. 8, Table 6), it is evident that reducing the concentration of circumferential fibers (i.e., increased dispersion) or replacing circumferential fibers with longitudinal fibers (three-layer model) can alter the degree of strain on the internal os of the cervix. This highlights the role of the cervix as a sphincter, particularly its ability to reconfigure its circumferential fibers to reduce strain under high loads without altering its stiffness [58]. The effect on cervical compression is reduced compared to that observed in variations in other material properties, thereby introducing cervical fiber structure or dispersion as a metric that is fairly independent of the previously mentioned cervical stiffness assessments. Currently, there are limitations in the overall quantification of cervical fiber organization, as well as a lack of methods and tools to routinely image the *in vivo* fiber structure of the cervix. The continued application of shear wave elastography may be promising in better characterizing physiological fiber orientations, as well as identifying fiber architectures that can be used as an additional marker to predict pathological labor conditions [93, 96, 97].

Lastly, although alterations in the fiber properties (*a*_*f*_, *κ*) in the uterus resulted in minimal changes in uterine tensile strain, increasing *E* in the uterus led to a notable reduction in tensile strain. Uterine distension has been found to modulate the excitability of the uterus through inflammation and stretch-mediated excitation and contraction [16–18]. Excessive early uterine distension has been found to be a potential pathway for preterm birth as early inflammation and activation of stretch-mediated excitation pathways prematurely drive the uterus into an excitable state[17, 18]. Thus, such investigations will become increasingly relevant upon implementation of a stretch-mediated active contraction model [9, 98]. As a step toward this direction, we have developed, in parallel, a multiscale framework to model myometrial activation and action potential propagation during late gestation and applied it to understand the role of ion channels and gap junctions in regulating uterine excitability [14]. Future work will couple the current mechanics model with the conduction model to simulate biophysically informed contraction patterns.

We acknowledge the limitations of this study. The present study focuses on the development of the model configuration methods; however, varying levels of fidelity can continue to be incorporated based on the needs of future studies of the physiological response of the uterus and cervix.

First, our current methodology has been developed for a single-patient geometry using a simplified, parameterized model, although it is derived from patient images. Future studies will focus on translating model parameters into a larger cohort of patient-specific models to optimize boundary condition parameter selection methods and investigate the effects of varying levels of geometric fidelity. Specifically, the study by Louwagie et al. discusses improved-fidelity parametric approaches for constructing uterine geometry with fewer simplifications of the coronal plane profile [22], which will be considered for future work.

Second, although widely followed [19, 21], our work assumes that the uterus is in a stress-free state (i.e., the reference configuration) at the beginning of loading, with subsequent application of IUP. However, because the patient images were captured during pregnancy, the geometry is preloaded with IUP, which limits the physiological relevance of the assumed loading state. As we currently lack imaging of the gravid uterus in a stress-free state, we look forward to addressing this limitation through future growth and remodeling studies or by prestressing. Despite this limitation, we believe that the comparative trends identified in the sensitivity analysis would still hold when accounting for a reference configuration. Although strain magnitudes will differ, the mechanistic role of the fibers and ground substance will likely remain consistent, maintaining patterns found here relative to the baseline configuration. We also assumed that the applied load is uniformly distributed across the entire inner surface of the uterus and cervix. However, fetal movements through the amniotic fluid and the presence of the placenta could introduce non-uniform hydrodynamic pressure and stress concentrations that are not accounted for in our model.

Third, the largest difference in deformation was observed between the two modeling approaches in the lateral region of the geometry. To minimize the number of tuned parameters and the cost of manual tuning, the current Robin boundary condition configuration prioritizes matching displacements in the anterior-posterior and distal-medial directions. Additionally, the Robin boundary conditions used here employ omnidirectional springs at each node, limiting their ability to capture a sliding interface between the uterus and abdomen, which is representative of the lubricating effects of the perimetrium. Within the cardiac mechanics field, the sliding effect representative of the pericardium has been implemented through reduced stiffnesses in the tangential directions [36]. To efficiently address both issues, future model development approaches could explore automated tuning pipelines to account for the increased number of parameters associated with spatially varying boundary conditions in both the normal and tangential directions.

Fourth, detailed fiber direction data is limited for the pregnant uterus and cervix. Currently, our model incorporates a single fiber family for both tissues with dispersion representative of experimental data. However, both tissues are highly heterogeneous and may be better represented with multiple fiber families that vary spatially within the tissue [46, 57, 89, 99]. To some extent, the fiber heterogeneity was evaluated by testing the three-layer fiber architecture of the cervix. However, this model is not comprehensive and still lacks spatial variation in fiber architecture across the cervix. In experimental mouse models, prior work has found greater organization representative of this three-layer model in the upper cervix and less prominent organization in the lower cervix [89]. Future work will continue to refine the current model through data collection and analysis of fibers from *ex vivo* uterine and cervical tissue samples, thereby further informing the fiber architecture model and its evolution throughout gestation.

Lastly, microstructurally-detailed material models have been proposed and fit to capture the mechanical properties of the uterus and cervix [66, 67, 100]. The GOH model, although simplified compared to these models, sufficiently captures the salient features of the mechanical response (anisotropy, hyperelasticity, and tension-compression asymmetry) while being computationally efficient and readily accessible. To continue investigating the physiological response of the uterus and cervix under loads and pathological conditions, these microstructure-based mechanical models may be used to provide greater fidelity in the tissue response.

In conclusion, we present a novel workflow for modeling passive uterine mechanics during late pregnancy that balances between fidelity and efficiency by leveraging robust cardiac mechanics modeling techniques. This approach lays the groundwork for future investigations into more complex myometrial excitation and contraction, as well as longitudinal studies of uterine mechanics during gestation.

## Abbreviations

EDD: Estimated Delivery Date
FEA: Finite Element Analysis
IUP: Intrauterine Pressure
LDRB: Laplace-Dirichlet Rule-Based
MRI: Magnetic Resonance Imaging
OCT: Optical Coherence Tomography

## Declarations

### Ethics Statement

The authors declare that they have no known competing financial interests or personal relationships that could have appeared to influence the work reported in this paper.

### Human Subjects

Patient-specific dimensions for model parameterization were taken from a publicly available dataset [101].

### Consent for Publication

All authors have agreed with the content and given explicit consent to submit the manuscript for publication.

### Data Availability

Data will be shared upon a reasonable request from the corresponding authors.

### Competing interests

The authors have no competing interests to declare.

### Funding Sources

NSF Graduate Research Fellowship (GRF); AHA Second Century Early Faculty Independence Award (24SCEFIA1260268); US NSF CAREER Award (2443726);

## Acknowledgments

The authors would like to acknowledge helpful discussions with Prof. Gerard Ateshian regarding FEBio model configuration and simulation setup. OM would like to acknowledge a graduate research fellowship through the US National Science Foundation. VV would like to acknowledge partial funding support from early career development awards, including the American Heart Association’s Second Century Early Faculty Independence Award (24SCEFIA1260268) and the US National Science Foundation’s CAREER Award (2443726). We also acknowledge computing resources from ACCESS (SDSC Expanse) and the Columbia Ginsburg HPC cluster for supporting this work.

## 5 Author Contributions

### Olivia Mergler

Data curation; Formal analysis; Investigation; Methodology; Software; Validation; Visualization; Writing – original draft; Writing – review & editing;

### Abigail Laughlin

Formal analysis; Investigation; Methodology; Writing – review & editing;

### Erin Louwagie

Formal analysis; Investigation; Writing – review & editing;

### Lei Shi

Formal analysis; Investigation; Methodology; Software; Writing – review & editing;

### Kristin Myers

Conceptualization; Formal analysis; Funding acquisition; Investigation; Methodology; Project administration; Resources; Supervision; Writing – review & editing;

### Vijay Vedula

Conceptualization; Formal analysis; Funding acquisition; Investigation; Methodology; Project administration; Resources; Supervision; Writing – review & editing;

## Appendix

**Table A1.**
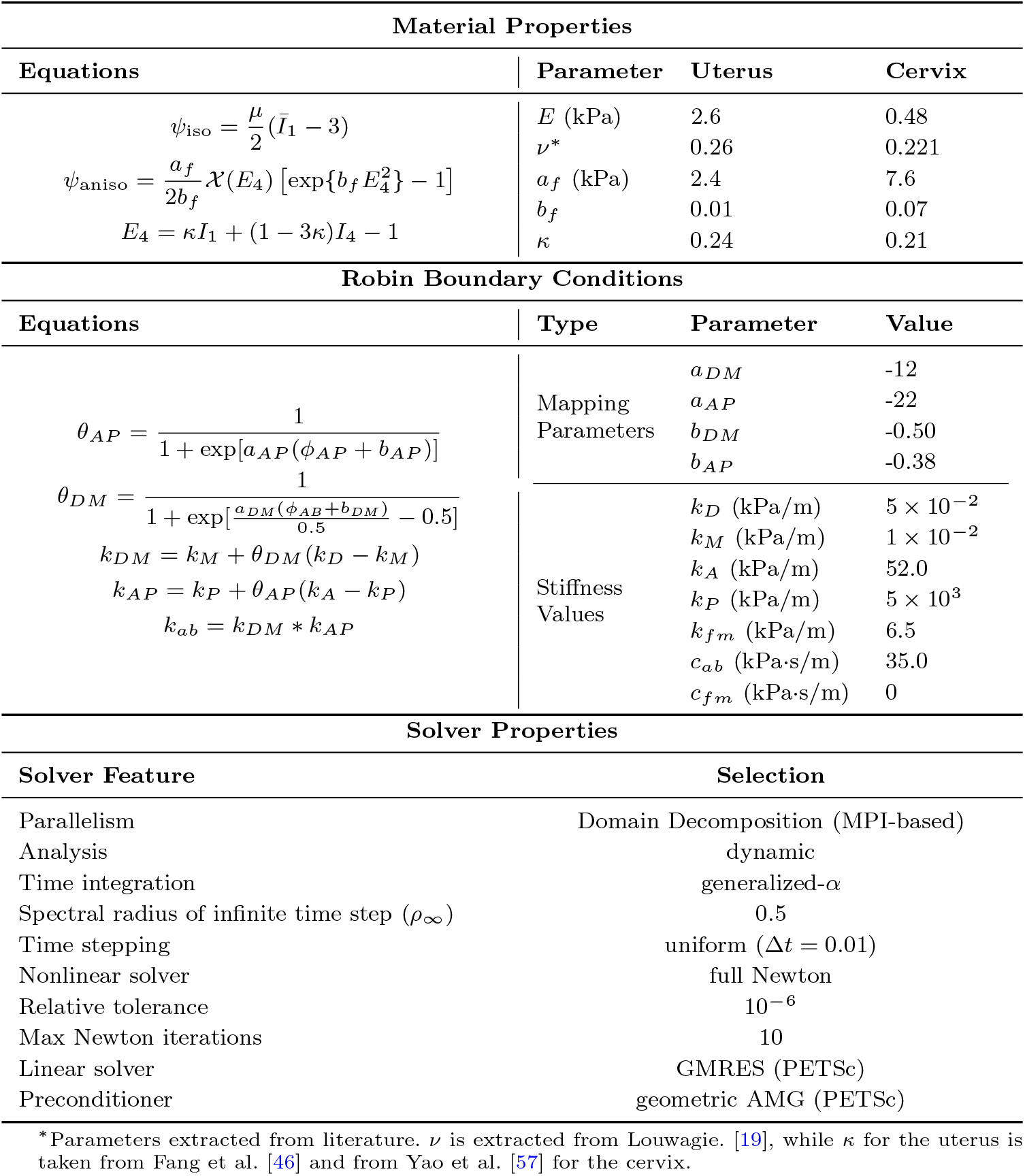
Summarized parameters and the corresponding equations for baseline model configuration.

**Table A2.** Nodal differences in Lagrange strain components between the two modeling approaches. Values reported as regional mean *±* standard deviation.

| Tissue Domain | Region* | Err$_{E_{xx}}$ | Err$_{E_{yy}}$ | Err$_{E_{zz}}$ | Err$_{E_{xy}}$ | Err$_{E_{xz}}$ | Err$_{E_{yz}}$ |
| --- | --- | --- | --- | --- | --- | --- | --- |
| Uterus | Overall | 0.029 ± 0.018 | 0.028 ± 0.029 | 0.026 ± 0.018 | 0.017 ± 0.015 | 0.013 ± 0.011 | 0.014 ± 0.016 |
| | AM | 0.032 ± 0.006 | 0.017 ± 0.003 | 0.018 ± 0.008 | 0.008 ± 0.005 | 0.007 ± 0.007 | 0.010 ± 0.007 |
| | AL | 0.027 ± 0.015 | 0.021 ± 0.024 | 0.015 ± 0.010 | 0.016 ± 0.010 | 0.011 ± 0.009 | 0.020 ± 0.019 |
| | PM | 0.031 ± 0.022 | 0.019 ± 0.013 | 0.038 ± 0.018 | 0.017 ± 0.017 | 0.014 ± 0.011 | 0.009 ± 0.008 |
| | PL | 0.025 ± 0.016 | 0.063 ± 0.041 | 0.027 ± 0.019 | 0.026 ± 0.019 | 0.018 ± 0.013 | 0.018 ± 0.020 |
| Cervix (Cx) | | 0.021 ± 0.024 | 0.018 ± 0.020 | 0.024 ± 0.026 | 0.015 ± 0.014 | 0.015 ± 0.017 | 0.014 ± 0.015 |
| Combined Model | | 0.028 ± 0.019 | 0.027 ± 0.029 | 0.026 ± 0.019 | 0.017 ± 0.015 | 0.013 ± 0.012 | 0.014 ± 0.019 |

\*Please refer Fig. 2b for more details on uterus partitions.

**Fig. A1.**
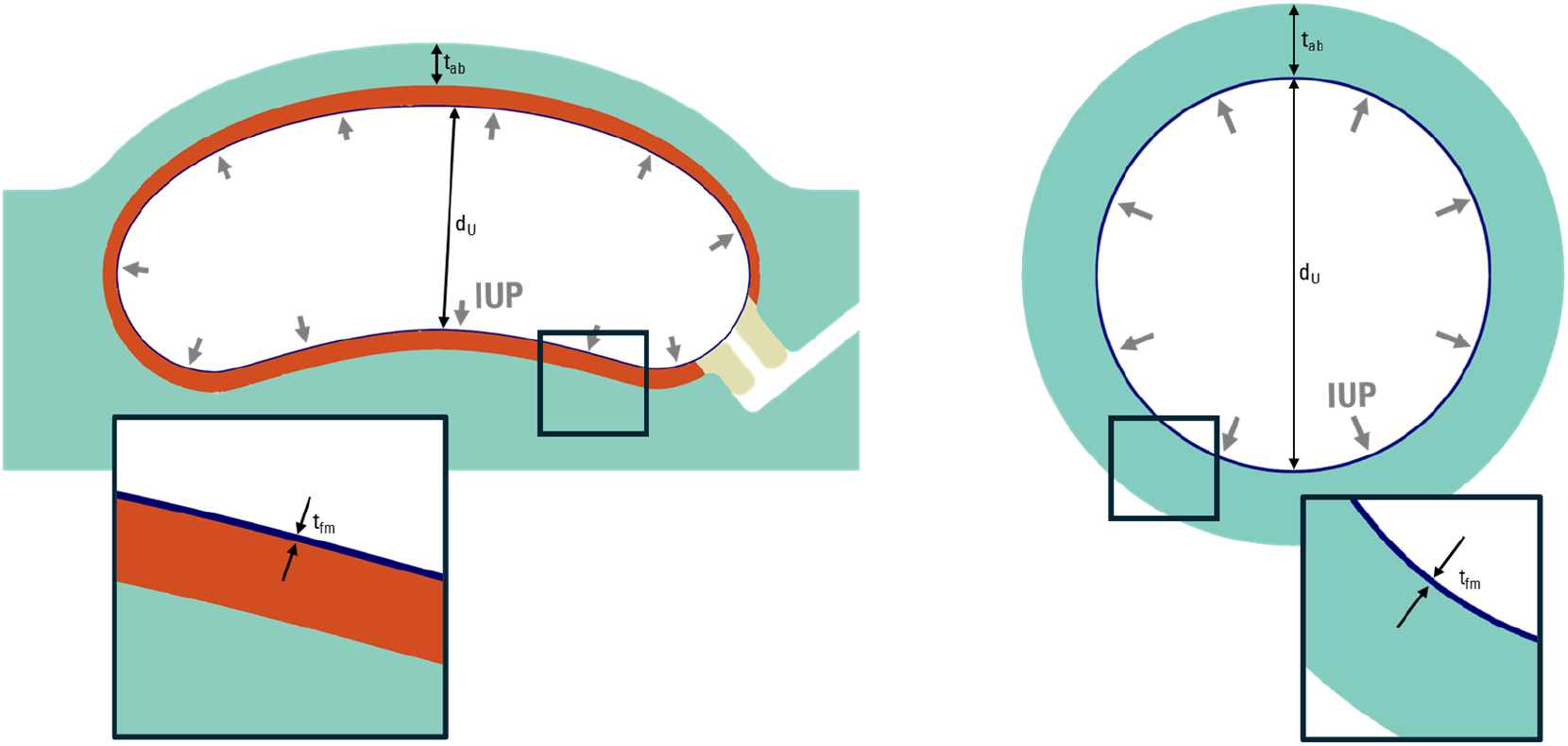
Configuration of spherical inflation tests for determining *k*_*fm*_ from *k*_*A*_. Only half a sphere is inflated with appropriate symmetry boundary conditions. Both half-spheres have an inner diameter consistent with that of the uterus (*d*_*U*_). The thickness of the abdomen (*t*_*ab*_) and fetal membrane *t*_*fm*_ spheres match those defined in the parameterized model. Spheres are overlaid for visualization; however, simulations for each half-sphere are computed independently, with no contact between them.

**Fig. A2.**
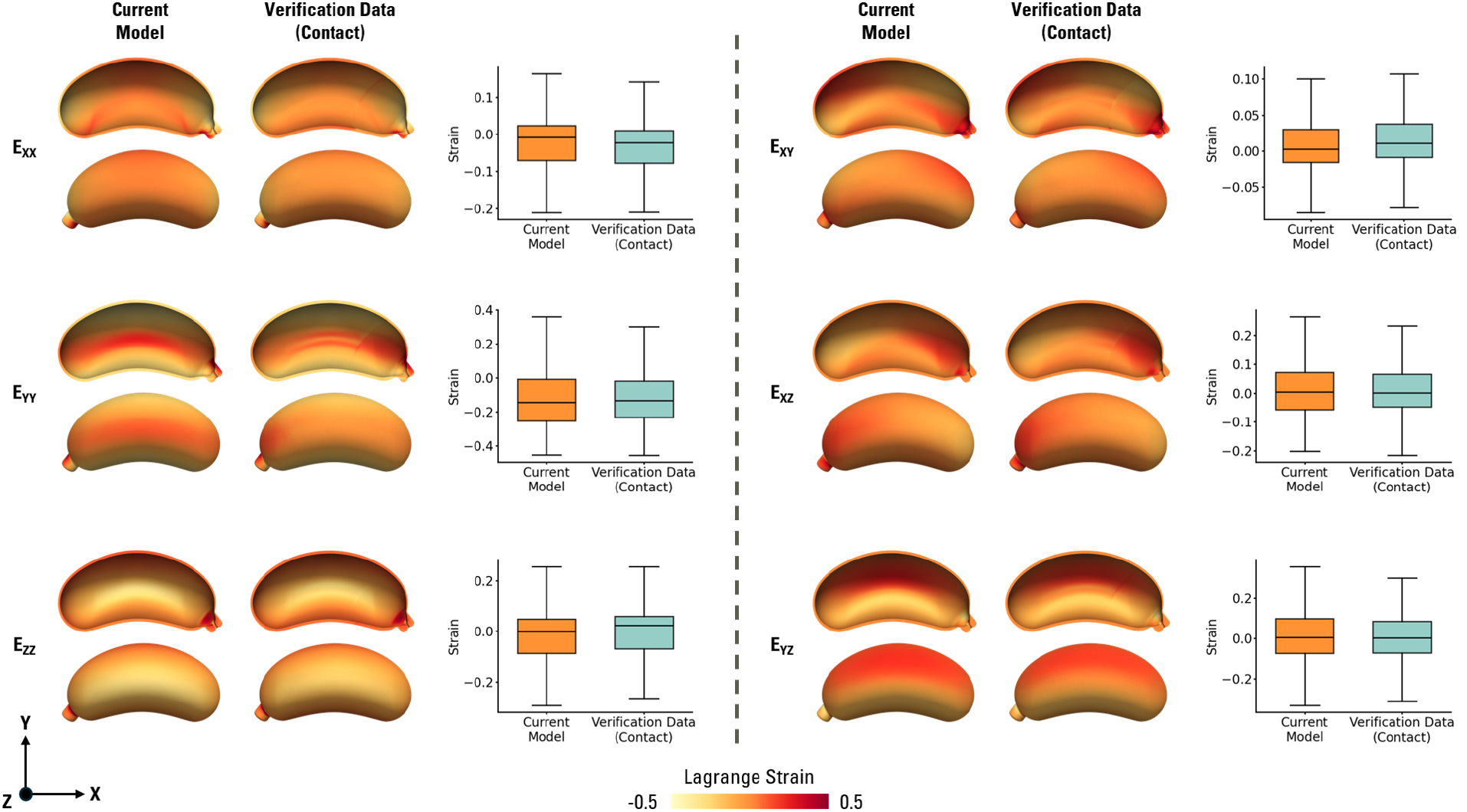
Comparison of Lagrange strain components between the current model and the verification data. For each strain component, the left two figures show the strain field for the current model configuration, and the center two figures show the strain field from the contact-based verification model. The plot to the right quantifies the strain distributions using a box-and-whisker plot.

## Footnotes

1 The myometrium is the middle layer of the uterus, mainly composed of smooth muscle cells, which are primarily responsible for inducing uterine contractions [15]

2 https://febio.org/

3 https://help.febio.org/docs/FEBioUser-3-6/UM36-4.1.4.11.html

4 https://github.com/SimVascular/svFSI

5 https://petsc.org/release/manualpages/PC/PCGAMG/

6 Portable, Extensible Toolkit for Scientific Computing (PETSc), version 3.18.5, https://petsc.org/release/

7 FEBio version 3.5.0, https://febio.org/

## Notes

### Competing Interest Statement

The authors have declared no competing interest.

